# Repeated blast mild traumatic brain injury and oxycodone self-administration produce interactive effects on neuroimaging outcomes

**DOI:** 10.1101/2020.11.18.388421

**Authors:** Matthew J. Muelbl, Breanna Glaeser, Alok S. Shah, Rachel Chiariello, Natalie N. Nawarawong, Brian D. Stemper, Matthew D. Budde, Christopher M. Olsen

## Abstract

Traumatic brain injury (TBI) and drug addiction are common comorbidities, but it is unknown if the neurological sequelae of TBI contribute to this relationship. We have previously reported elevated oxycodone seeking after drug self-administration in rats that received repeated blast TBI (rbTBI). TBI and exposure to drugs of abuse can each change structural and functional neuroimaging outcomes, but it is unknown if there are interactive effects of injury and drug exposure. To determine the effects of TBI and oxycodone exposure, we subjected rats to rbTBI and oxycodone self-administration and measured drug seeking and several neuroimaging measures. We found interactive effects of rbTBI and oxycodone on fractional anisotropy (FA) in the nucleus accumbens (NAc), and that FA in the medial prefrontal cortex (mPFC) was correlated with drug seeking. We also found an interactive effect of injury and drug on widespread functional connectivity and regional homogeneity of the BOLD response, and that interhemispheric functional connectivity in the infralimbic medial prefrontal cortex positively correlated with drug seeking. In conclusion, rbTBI and oxycodone self-administration had interactive effects on structural and functional MRI measures, and correlational effects were found between some of these measures and drug seeking. These data support the hypothesis that TBI and opioid exposure produce neuroadaptations that contribute to addiction liability.

## Introduction

Traumatic brain injury (TBI) and opioid abuse have each reached epidemic proportions. An estimated 2.4 million Americans suffer a TBI each year ^1^, and in 2017 there were over 47,000 deaths due to opioid overdose ^2^. The incidence of TBI among veterans of the military has been approximated at 30% ^3^, with blast injury causing the overwhelming majority of TBIs in combat settings. Co-morbidity between non-severe TBI and substance abuse has been found in numerous civilian and military epidemiological studies ^4–10^, and there is evidence that opioid abuse may be particularly prevalent following TBI ^11, 12^. For example, in a study of >40,000 Air Force personnel, those that sustained mild TBI (compared to non-brain injuries) had elevated hazard ratios for several drugs of abuse ^5^. Furthermore, the hazard ratio for opioids was higher than all other drugs of abuse in the 18 months following injury (including alcohol) ^5^. Despite extensive evidence of increased substance abuse disorder following TBI and high rates of opioid prescriptions to veterans and civilians that sustain TBI, little is known about how neurological damage associated with TBI may exacerbate opioid addiction liability.

We have previously demonstrated that our blast model of mild TBI leads to enduring microstructural changes in the medial prefrontal cortex ^13^, a brain region critically involved in drug seeking in humans and rodent models (reviewed in ^14–16^). We have also found that repeated blast mild TBI (rbTBI) persistently elevated levels of oxycodone seeking, despite similar levels of prior drug self-administration ^17^. In the current study, we hypothesized that rbTBI and oxycodone self-administration would lead to persistent changes in structure and/or function of the medial prefrontal cortex which may also extend to other aspects of mesocorticolimbic circuitry that would be associated with elevated drug seeking. To test this hypothesis, rats underwent rbTBI, oxycodone self-administration, and drug seeking sessions. Structural and functional MRI were performed at acute and chronic timepoints (1- and 7- weeks post-injury, respectively).

## Methods

An overview and timeline of experimental procedures is found in Fig 1A.

**Figure 1:**
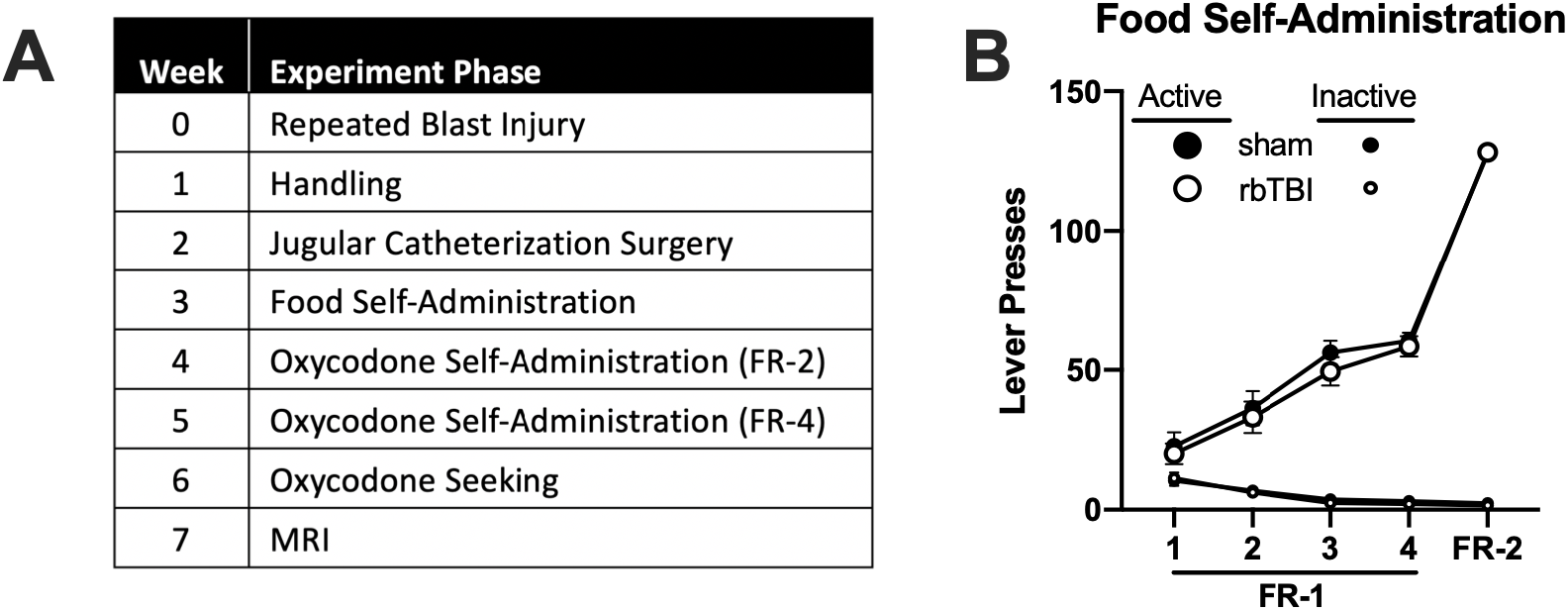
A) Experimental timeline. B) Acquisition of food self-administration in sham and rbTBI mice.

### Animals

Male Sprague Dawley rats (Harlan/Envigo, Madison, WI, USA) arrived at the Zablocki Veterans Affairs Medical Center (ZVAMC) and were housed in pairs. Three days after the final blast or sham exposure, rats were transferred from the ZVAMC to Medical College of Wisconsin (MCW, approximately 5 miles away). Upon arrival at MCW, rats were singly housed for the remainder of the study. Food and water were provided ad libitum throughout the study, except when rats were in the behavioral apparatus. Animals were housed under a reverse light cycle (lights off 0730–1930), and experiments were conducted during the dark period. Experiments were conducted 5 days/week by experimenters blinded to the treatment condition. All experiments were conducted in compliance with the Institutional Animal Care and Use Committee (IACUC) at MCW and ZVAMC (protocol number: AUA 2816).

### Repeated blast exposure

Rats were exposed to repeated blast overpressure or repeated sham conditions as described in ^13, 17–19^. During blast or sham treatment, rats were maintained under anesthesia with a continuous delivery of 1.5% isoflurane and administered Rimadyl (carprofen, 5 mg/kg, s.c., Zoetis Inc, Kalamazoo, MI, USA). The torso of the rat was shielded with a metal cylinder, earplugs were inserted, and the head was restrained to prevent injury from rotational acceleration ^13^. Blast– exposed rats were positioned 17 cm from the end of a custom shock tube (3.6 cm inner diameter, 3.0 m driven section and 0.3 m driver section with Mylar membrane between the driver and driven sections) and 18 deg off axis from the shock tube to prevent interaction of the head with exhaust gases. Blast overpressure (450 kPa, 80 kPa*ms) was produced by pressurizing the driver section with helium until the membrane ruptured. Sham–exposed rats were subjected to the same procedures but were placed outside the overpressure shockwave. Following exposure, rats were placed on a heating pad and observed until return of righting reflex. Time from initiation of anesthesia prior to blast exposure until return of the righting reflex was quantified as ‘recovery time’. Following recovery, rats were transferred back to their home cage, and periodically observed for 6 h. This procedure was conducted daily for 3 days as a repeated blast injury model. Sham–exposed rats received sham procedures daily for 3 days. Experimenters were blind to group assignment during subsequent phases of experiments.

### Jugular catheterization surgery

Jugular catheterization was performed as described ^17, 20, 21^. Rats were anesthetized with isoflurane (3–5% induction, 1–3% maintenance), and a silicone catheter (Silastic, ID: 0.51 mm, OD: 0.94 mm; Dow Corning, Auburn, MI, USA) was implanted into the right jugular vein and exited via the interscapular region. The catheter was connected to a cannula assembly: a 22– gauge stainless steel cannula (Plastics One, Roanoke, VA, USA) mounted on a base made from dental acrylic and nylon mesh, which was implanted subcutaneously. At the start of the surgery and the following day, Rimadyl (carprofen, 5 mg/kg, s.c. Zoetis Inc) was given. Catheters were flushed daily with 0.2 mL of heparinized saline (30 units/mL) and cefazolin (100 mg/mL). Rats recovered for at least 1 week prior to the start of the drug self–administration experiments.

### Food training

Food training and oxycodone self-administration were performed as described in ^17^. Food and oxycodone self-administration and the seeking test all took place in operant conditioning chambers (31.8 × 25.4 × 26.7 cm, Med Associates, Fairfax, VT, USA) which were enclosed in sound–attenuating boxes. Chambers were equipped with two retractable levers on either side of a central food receptacle on the right wall, cue lights above each lever and a house light located on the left wall near the ceiling. Computer-controlled infusion pumps held syringes that were connected to tubing and liquid swivels. Sound attenuating boxes had a software-controlled exhaust fan.

Rats began by learning to self-administer food (fruit punch flavored 45 mg sucrose tablets, LabDiet, St. Louis, MO, USA) without food restriction. Each session lasted for 2 h/day unless the maximum number of rewards (64 sucrose pellets) was met prior to the conclusion of the 2 h. At the beginning of each session, the fan was turned on, both levers were extended, and a sucrose pellet was delivered. Rats were first trained on a fixed ratio (FR)–1 schedule of reinforcement, with a single press on the active lever resulting in sucrose pellet delivery. After meeting criteria (three consecutive days with ≥ 50 rewards earned and ≥ 70% of total lever presses on the active lever, 4 day minimum), rats performed a single food self-administration session on an FR-2 schedule of reinforcement (2 active lever presses required per reward).

The active lever was assigned to be on either the left or right side, and lever assignment was counterbalanced between rats within each group. After reward delivery, a 10 s timeout began, and the cue light above the active lever was illuminated for 5 s. During the timeout period, active and inactive lever responses were recorded but had no programmed consequence. Throughout the entire session, responses on the inactive lever were recorded but also had no programmed consequence.

### Oxycodone self-administration and seeking

After completion of food training, rats in the rbTBI and sham groups were further divided into those that self-administered oxycodone or received yoked infusions of saline. Each yoked saline rat was matched to a “master” rat from the same cohort. When the “master” rat earned an infusion of oxycodone, the yoked rat received an infusion of saline at the exact same time (computer controlled). Thus, the infusion history between the “master” and “yoke” rats is identical, controlling for the potential influence of infusions. Yoked saline rats still received cue light presentation in response to completing the response requirements. The infusion volume and time varied for each rat to ensure the appropriate dose was delivered from the stock solution (e.g., a 300 g rat received a 40 μL infusion over 1.6 s) and yoked saline rats received weight-corrected volumes of saline. This phase consisted of 10 daily 2–h sessions (maximum 96 infusions/session): 5 d each on FR-2 and FR-4 schedules of reinforcement. Catheter patency was determined after completion of the self–administration sessions using Brevital (methohexital, 9 mg/kg, i.v.; JHP Pharmaceuticals, Rochester, MI, USA) and any rat that did not meet criteria for patency (sedation within 10 s) was removed from the study. Three days after completion of the 10-day oxycodone self-administration (or yoked saline) phase, all rats were placed back into the operant conditioning chambers for a drug seeking test under extinction conditions. Completion of the FR-4 schedule resulted in saline infusion and cue delivery for all animals (i.e., yoked saline animals were no longer yoked).

### Magnetic Resonance Imaging (MRI)

In vivo magnetic resonance imaging was performed on a Bruker 9.4 T Biospec System. A Bruker Biospin 4-channel surface coil array was used for signal reception and transmission was achieved via a 72 mm diameter quadrature volume coil. Heart rate (ECG), core temperature, and respiration were monitored continuously. Anesthesia was induced with isoflurane (3.0%), which was tapered to 0% after initiation of constant tail vain infusion of dexmedetomidine. Dexmedetomidine was delivered via constant tail vain infusion (initial concentration: 50 μg/kg/hr) and was titrated to maintain anesthetic depth as determined by ECG, body temperature, and respiratory rate. Anesthetized rats were secured to a custom-made temperature-controlled cradle to minimize motion. A high-resolution anatomical image was acquired with an isotropic resolution of 134 μm^3^. For diffusion imaging, a 4-shot spin-echo echo planar imaging (DW-EPI) sequence was used as described previously ^22^ with TR=2000 and TE=23 ms, including fat saturation using both chemical shift-selective saturation and gradient reversal of the refocusing pulse slice. Thirty slices covered the entire brain at an in-plane resolution of 0.30×0.30 mm^2^, slice thickness of 0.8 mm and 0.3 mm slice gap. Two signal averages were used to acquire 60 unique diffusion directions ^23^ at each b-value of 250, 1000, and 2000 s/mm^2^ with 8 non-diffusion-weighted images. For resting state functional MRI, 200 images were acquired at a temporal resolution of 3 sec using single-shot gradient echo EPI at the identical spatial resolution. Three images were acquired for each repetition at echo times of 16.5, 33.1, and 49.6 ms, and each run was repeated 3 times for a total of 30 minutes for fMRI data acquisition. To examine for any evidence of overt injury or hemorrhage, a 3D multiple echo acquisition with 0.175 mm^3^ isotropic voxels (TR=50 ms) was acquired with 10 echo times between 2.2 and 24.9 ms. Across all animals, no hemorrhage was evident based on visual inspection.

### Statistical Analysis

Statistical tests were performed using Prism 8 (GraphPad, San Diego, CA, USA) or SPSS Statistics 24 (IBM, Inc.) software. Data were analyzed using two-way mixed model ANOVAs (session: within-subjects factors, experimental group: between-group factor) with Holm-Sidak post hoc tests. ANOVAs were performed using ranked data if data did not fit a normal distribution as determined by D’Agonstino & Pearson tests. Correlational analyses were performed using linear regressions (Spearman rank correlations if data did not fit a normal distribution) followed F tests to determine if slopes were significantly different from zero. Differences were considered significant when p ≤ 0.05.

## Results

### Behavioral measures

Repeated blast TBI resulted in shorter recovery time (36.6 ± 8.1 sec faster) from anesthesia compared to sham treated rats (F(1, 38)=20.26, η^2^=0.20 p<0.0001), but there was no effect of blast exposure number or interaction of blast with exposure number (both η^2^<0.10, p>0.39). On week three following rbTBI or sham treatment, rats underwent food self-administration training. Food self-administration did not differ between rats exposed to rbTBI and sham (main and interaction effects of rbTBI all η^2^<0.045, p>0.39; Fig 1B). These data indicate that there was no impact of injury on the ability to acquire and perform this task, a critical consideration in TBI research that utilizes operant conditioning ^24^. Next, rats either self-administered oxycodone or received yoked infusions of saline. Specifically, rats were split into pairs: when the “master” rat earned an infusion of oxycodone, this triggered a time-matched infusion of saline into the “yoked” rat. There was no significant difference between rbTBI and sham rats in oxycodone self-administration as measured by active lever presses during each FR phase (main and interaction effects of rbTBI all η^2^<0.065, p>0.21; Fig 2A) or number of infusions earned during the FR-2 phase (main and interaction effects of rbTBI both η^2^<0.045, p>0.26; Fig 2C) or the FR-4 phase (main and interaction effects of rbTBI all η^2^<0.083, p>0.16; Fig 2D). There was also no significant difference in lever pressing between rbTBI and sham rats in the yoked saline groups (main and interaction effects of rbTBI both η^2^<0.016, p>0.67; Fig 2B). This demonstrates that rbTBI does not lead to generalized deficits in extinction learning (i.e., extinction of sucrose pellet seeking); a significant finding considering that we have previously reported that rbTBI resulted in elevated oxycodone seeking in extinction conditions ^17^ and several studies have found that mild TBI impaired extinction of conditioned fear ^25–28^. Since we had previously demonstrated a TBI-associated increase in drug seeking using the same TBI and oxycodone self-administration procedures ^29^, we performed a planned comparison on oxycodone seeking between sham and rbTBI animals. Compared to sham injured rats, rbTBI rats had greater drug seeking during testing under extinction conditions (no drug available) (t(10)=2.38, d=0.72, p=0.039; Fig 2E).

**Figure 2:**
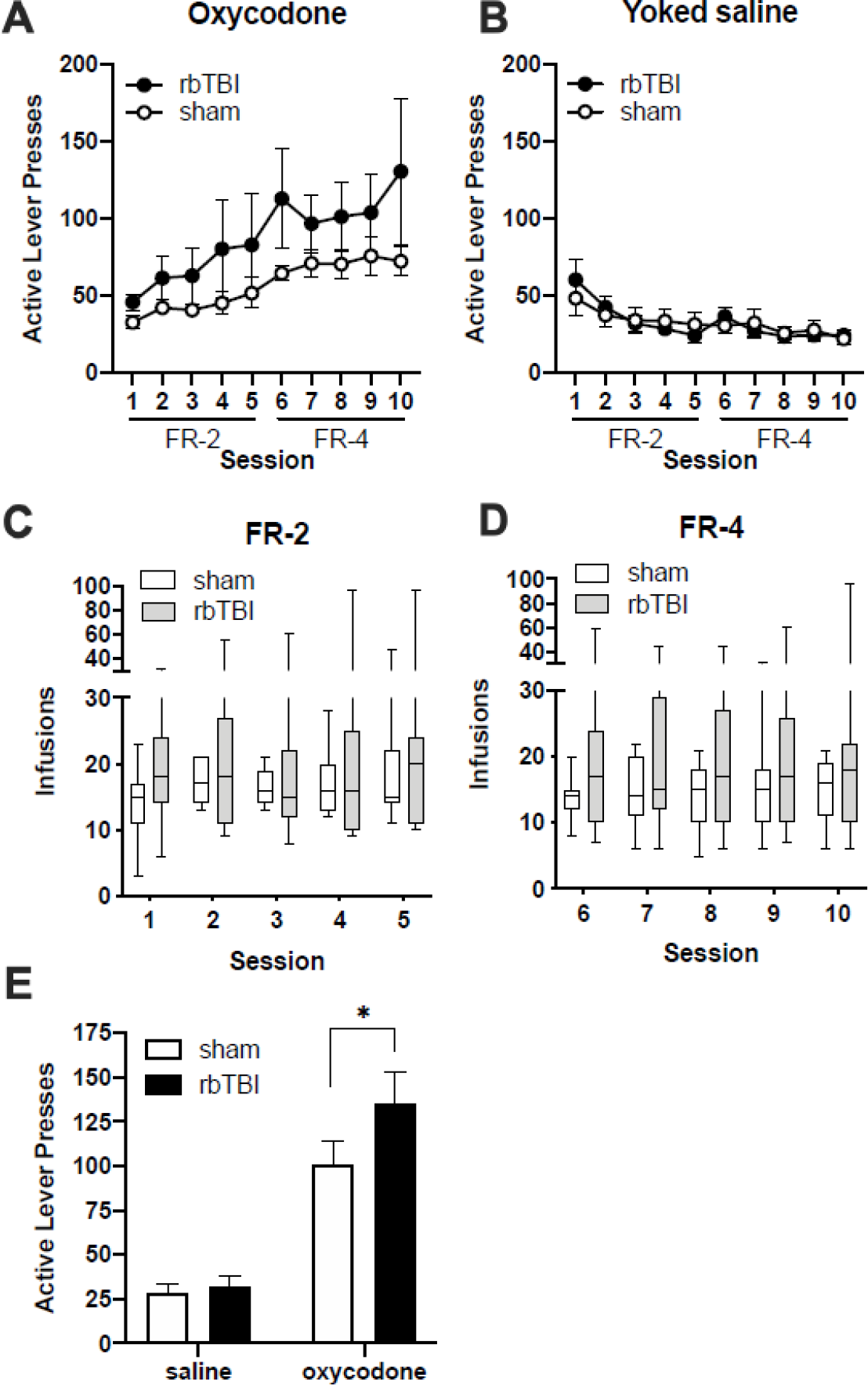
Oxycodone self-administration and seeking. A) Active lever responses for oxycodone. B) Active lever responses in rats that received yoked saline infusions. C, D) Oxycodone infusions earned during FR-2 (C) and FR-4 (D) sessions. E) Active presses during extinction trial. Boxes represent median and quartiles, whiskers represent range. Bars represent mean ± SEM. *p≤0.05

### MRI

To examine neuroimaging correlates of injury and oxycodone self-administration, rats underwent structural and functional MRI at 7-weeks post-injury, approximately 10 days following cessation of oxycodone self-administration.

#### Structural MRI

Structural assessment was performed by examining T2, T2*3D, fractional anisotropy (FA), and mean diffusivity (MD).

#### Factorial analysis of fractional anisotropy and mean diffusivity

We examined main and interactive effects of rbTBI and oxycodone self-administration on microstructure of the prefrontal cortex and nucleus accumbens. These brain regions are engaged during drug craving in human opioid users and drug seeking behaviors in rodent models of opioid addiction ^15, 30, 31^. These areas are also subject to damage in human blast and non-blast mild TBI ^32–35^. There were no significant main effects of injury or oxycodone, but a significant interaction was found in the FA of both hemispheres of the nucleus accumbens (p≤0.05).

#### Relationship between fractional anisotropy and drug seeking

We have previously demonstrated that bTBI lead to changes in hippocampal FA that correlated with spatial learning deficits ^13^. To examine the possibility that mPFC FA was associated with drug seeking, we performed correlational analyses between FA in the subregions of the prefrontal cortex (PFC) and nucleus accumbens (NAc) and active lever presses during the drug seeking session. We examined sham and rbTBI rats separately and together to determine if associations were specific to injured rats or if associations generalized to both populations. Significant associations were only found in the subregions of the mPFC ipsilateral to injury (Fig 3). In the prelimbic PFC (PrL), there were significant linear correlations in the rbTBI group (r^2^=41, p=0.047) and across injury groups (r^2^=23, p=0.020), but not in sham injured rats (r^2^=11, p=0.37). In the infralimbic PFC (IL), there was only a significant association between FA and drug seeking when the sham and injury groups were combined (r^2^=22, p=0.046). To determine if this association was explained by the amount of oxycodone intake during prior self-administration sessions, we examined the correlations between the PFC/NAc and drug intake (Table 1). There was no association between either ipsilateral subregion and drug intake (Pearson’s r and Spearman’s rho all <0.48), however, there was a positive linear association between the contralateral IL and prior oxycodone intake in rbTBI (rho=0.65, p=0.049) and in combined sham and injury rats (rho=0.59, p=0.0084). These data suggest that the association between reduced FA ipsilateral to injury and elevated drug seeking is not due to the amount of oxycodone self-administered, and that there is a positive relationship between total oxycodone intake and FA in the contralateral IL mPFC.

**Table 1:** Correlations between structural neuroimaging measures and seeking or intake. FA: fractional anisotropy, MD: mean diffusivity. Ipsilateral: left, contralateral: right. Italicized values represent Spearman’s rho instead of Pearson’s r. *p≤0.05 **p≤0.01.

| FA v. Seeking |  | sham | rbTBI | sham+rbTBI |
| --- | --- | --- | --- | --- |
| Ipsilateral | PrL | -0.34 | **-0.64\*** | **-0.53\*** |
|  | IL | -0.46 | -0.46 | **-0.46\*** |
|  | NAc | 0.30 | -0.42 | -0.19 |
| Contralateral | PrL | 0.10 | -0.03 | -0.03 |
|  | IL | 0.35 | 0.30 | 0.30 |
|  | NAc | -0.26 | -0.37 | -0.26 |

| MD v. Seeking |  | sham | rbTBI | sham+rbTBI |
| --- | --- | --- | --- | --- |
| Ipsilateral | PrL | 0.42 | -0.05 | 0.12 |
|  | IL | 0.25 | -0.12 | 0.02 |
|  | NAc | 0.10 | -0.12 | -0.01 |
| Contralateral | PrL | 0.03 | -0.07 | 0.11 |
|  | IL | -0.10 | -0.36 | -0.16 |
|  | NAc | 0.26 | 0.01 | 0.05 |

| FA v. Intake |  | sham | rbTBI | sham+rbTBI |
| --- | --- | --- | --- | --- |
| Ipsilateral | PrL | -0.03 | -0.32 | -0.31 |
|  | IL | -0.48 | -0.36 | -0.36 |
|  | NAc | 0.48 | 0.04 | 0.03 |
| Contralateral | PrL | 0.15 | 0.05 | 0.16 |
|  | IL | 0.39 | **0.65\*** | **0.59\*\*** |
|  | NAc | 0.11 | 0.07 | 0.01 |

| MD v. Intake |  | sham | rbTBI | sham+rbTBI |
| --- | --- | --- | --- | --- |
| Ipsilateral | PrL | -0.03 | -0.30 | -0.19 |
|  | IL | -0.10 | **-0.67\*** | -0.36 |
|  | NAc | -0.27 | -0.40 | -0.31 |
| Contralateral | PrL | -0.32 | -0.29 | -0.17 |
|  | IL | -0.15 | **-0.67\*** | **-0.55\*** |
|  | NAc | -0.15 | -0.52 | -0.30 |

**Figure 3:**
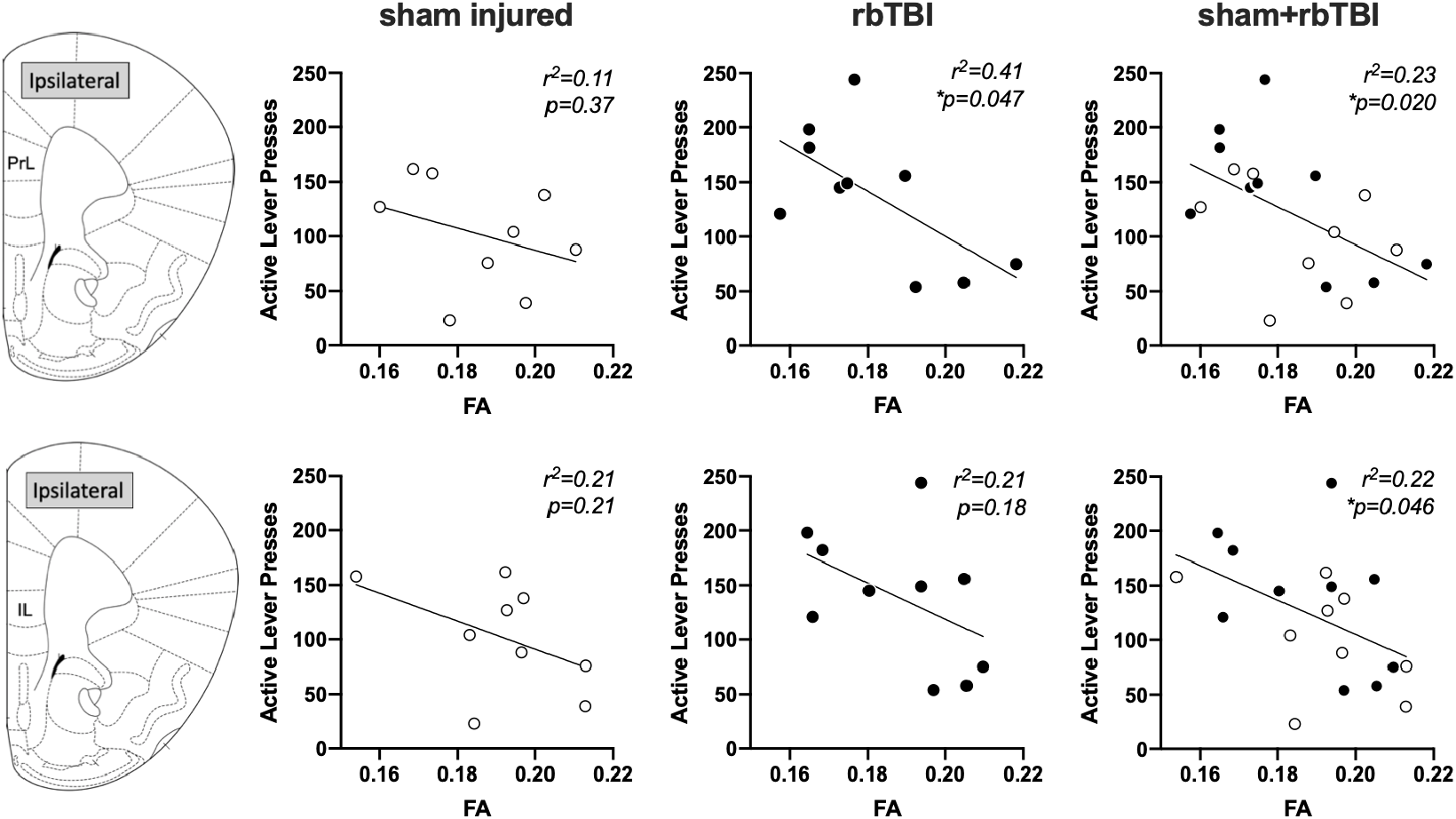
Correlations between FA and active lever presses in ipsilateral PrL and IL. *p≤0.05

##### Relationship between mean diffusivity and drug seeking

We examined mean diffusivity (MD) to determine if this additional measure of local microstructure was associated with oxycodone seeking or intake during self-administration (Table 1). No significant associations were found between MD and drug seeking in any region, although the MD in both hemispheres of the IL mPFC had a significant negative association with oxycodone intake in rbTBI rats (rho=−0.67, p=0.039 for each hemisphere). Similarly, there was a significant negative association between MD and oxycodone intake when injury groups were combined (rho=−0.55, p=0.014).

##### Functional MRI

###### Regional homogeneity (REHO)

**Figure 4:**
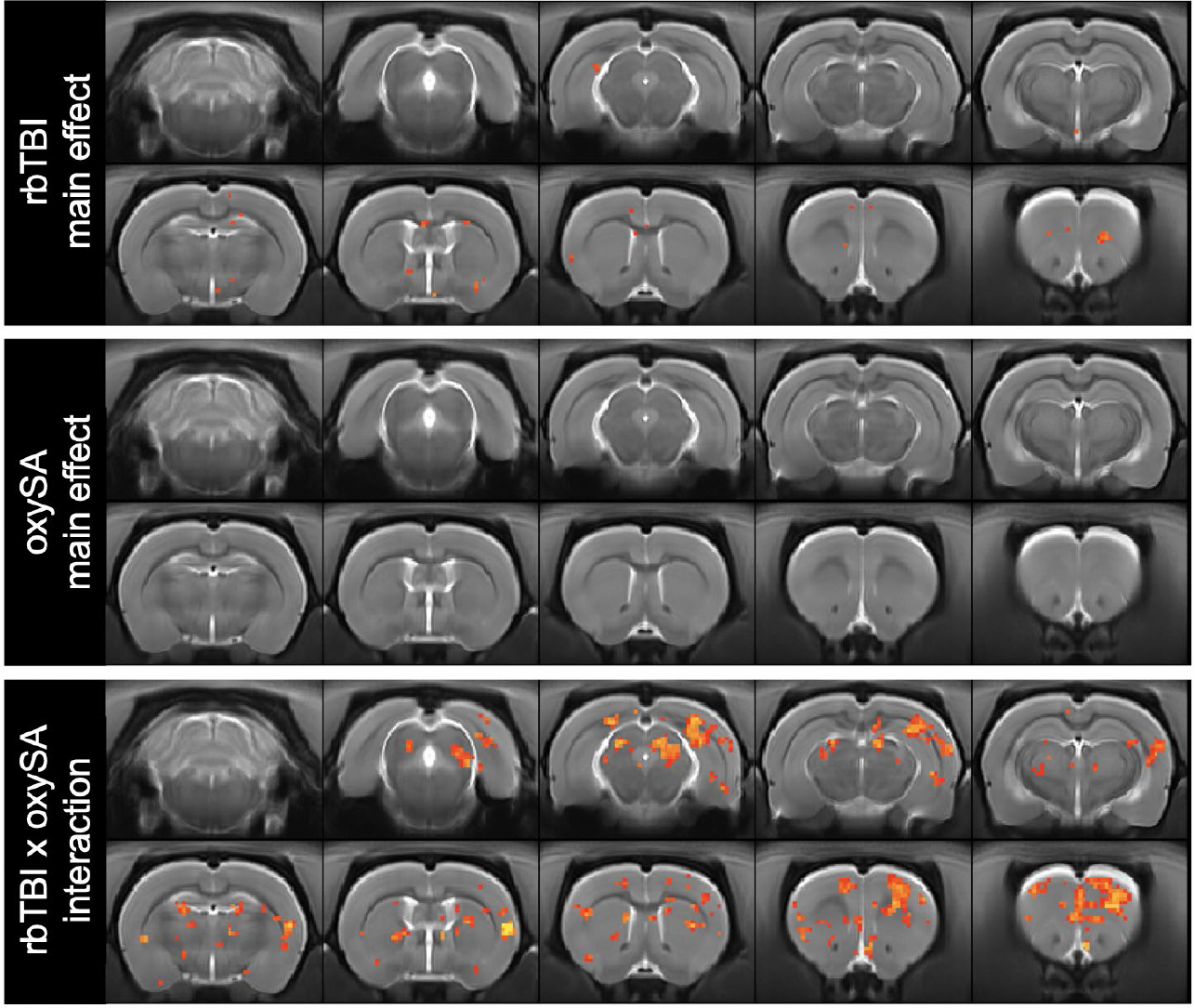
Regional homogeneity (REHO) of BOLD signal. Red/yellow voxels represent main or interaction effects at p≤0.05.

##### Global functional connectivity

Regions of interest were represented as nodes and edges were calculated as the temporal Pearson correlation coefficient in the BOLD response between pairwise nodes for each of the four treatment groups (Fig 7). rbTBI/saline and sham/oxycodone self-administration rats showed comparable patterns of connectivity to sham/saline rats, while rbTBI/oxycodone self-administration rats had a larger number of connected nodes, as defined by Pearson r≥0.6 (Fig 7). Edge strengths were tested in a factorial design, which also yielded a significant interaction (p≤0.05), but no significant main effects (both p>0.62).

**Figure 5:**
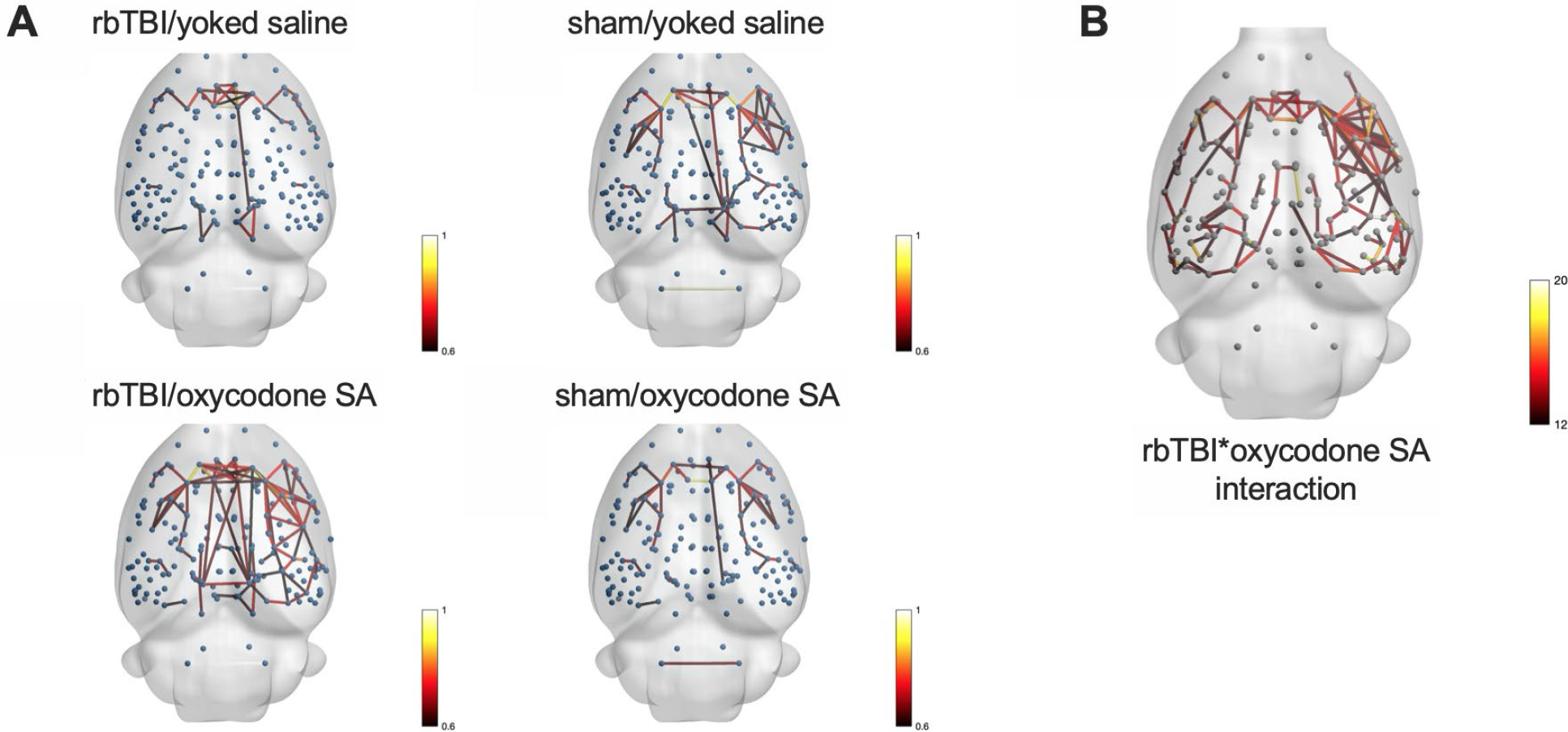
Network-based statistics. A) Each node (blue sphere) represents a region of interest, each edge represents a significant correlation between nodes. B) Nodes connected by edges had a significant interaction between rbTBI and oxycodone self-administration. Edges are colored based on Pearson r values (A) or F values (B).

#### Relationship between functional connectivity and drug seeking

Prefrontal-accumbens neurotransmission has been implicated in drug seeking for several drugs of abuse, including opioids (reviewed in ^15, 36, 37^). We tested the hypothesis that functional connectivity between regions of the PFC and nucleus accumbens would be positively correlated with oxycodone seeking, and this would be relationship would be independent of injury. Pairwise correlations were performed on each hemisphere of the PrL, IL, and NAc. Analysis of the relationship between functional connectivity and drug seeking (active lever presses during extinction session) found that only the inter-hemispheric connectivity of the IL was significantly correlated with drug seeking in injured and uninjured rats (Fig 6C). Prefrontal-accumbens functional connectivity was significantly associated with drug intake on the left hemisphere (ipsilateral to injury in rbTBI rats), and this was observed in rbTBI and the combined population of rats (Fig 6E,F).

**Figure 6:**
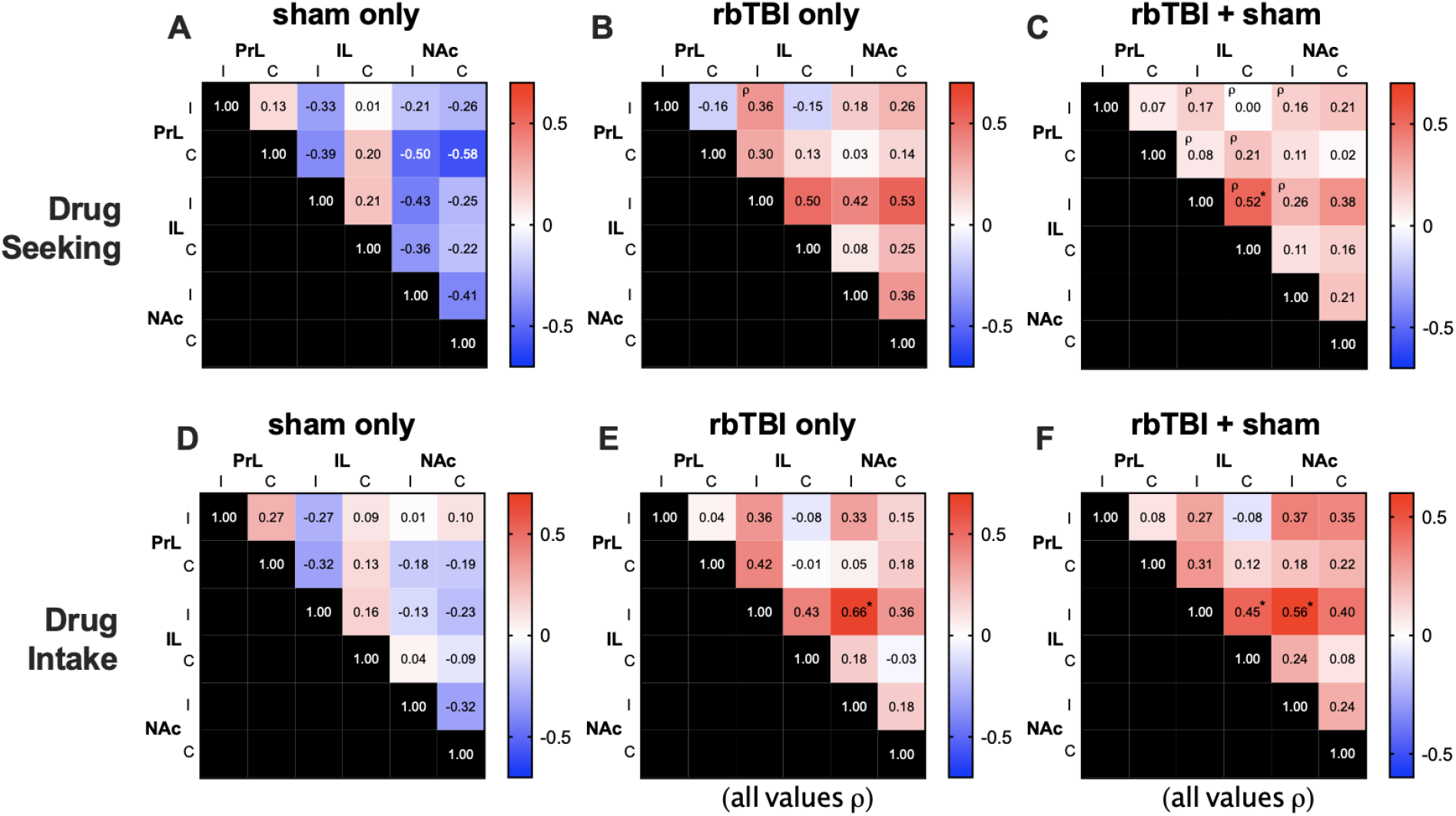
Relationship between functional connectivity and drug seeking (A-C) or drug intake (D-F). Values are Pearson r except where denoted by ρ. Significant correlations are identified in bold text. *p≤0.05. I, C: ispi- or contra-lateral to injury, corresponding to left or right hemisphere, respectively in all rats.

## Discussion

We have previously shown that rbTBI-exposed rats had elevated drug seeking following oxycodone self-administration despite similar levels of drug intake ^17^. In this study, we used neuroimaging to explored potential interactive effects and correlates of behavior on structural and functional brain networks. We focused on the prefrontal cortex and nucleus accumbens due to their known involvement in persistent drug seeking behavior ^15, 38, 39^. We found several neuroimaging metrics that were associated with drug seeking behavior, prior drug intake, and interactive effects of injury and oxycodone self-administration. Microstructural analysis identified significant negative associations between FA and drug seeking in the left (but not right) PrL and IL PFC, the side ipsilateral to injury in rbTBI rats. There was also a significant positive association between FA and drug intake during self-administration in the right hemisphere, while the association between MD and prior oxycodone intake was bilateral. The effect was largest in rbTBI rats but was also significant in the combined population of injured and un-injured rats. Within the NAc, there was a significant interaction between injury and oxycodone self-administration on FA, but neither FA nor MD were associated with the amount of oxycodone self-administered or subsequent drug seeking. All of the microstructural correlations with behavior had larger effects in injured relative to uninjured rats, suggesting that rbTBI enabled or enhanced the emergence of a relationship between structural parameters of the PFC and prior drug seeking and/or drug intake. Resting state functional connectivity (rsfc) analyses found a widespread increase in network connectivity only in rats that received injury and oxycodone SA. There were also significant correlations between IL intra-hemispheric connectivity and drug seeking, and between ipsilateral IL – NAc connectivity and oxycodone intake during self-administration. As observed in the structural measures, these effects were largest in injured animals, suggesting that similar to our structural outcomes, the interaction of injury and oxycodone exposure enabled the emergence of a relationship between functional connectivity and prior drug seeking and/or drug intake. An interaction was also measured in local connectivity: regional homogeneity was minimally affected by injury or oxycodone self-administration alone, but animals receiving both had marked increases in voxelwise homogeneity of the medial PFC, NAc, and other areas, such as motor cortex, hippocampus, deep layers of the superior colliculus, and external cortex of the inferior colliculus.

The present study adds to a growing body of preclinical work that has largely demonstrated that experimental TBI is associated with worse outcomes in models of substance abuse. For example, studies across different species and TBI models have found elevated alcohol intake following injury ^18, 40–43^(although see ^44^). Experimental TBI has also been shown to increase cocaine conditioned reward ^45, 46^ and self-administration ^47^ (although see ^20^). In the same model of repeated blast mild TBI used here, we found that injured animals had reduced oxycodone self-administration, but drug seeking was elevated during extinction testing ^17^. Thus, consistent with clinical studies that report associations between TBI and substance abuse ^4–12^, there is a growing body of preclinical work that supports a causative role for TBI in elevated addiction liability. In the current study, we found that drug seeking was elevated after injury and oxycodone SA. We also found structural and functional changes in the brain, with some of these alterations associated with prior drug seeking or intake.

The significance of injury and opioid exposure has been examined in peripheral and central injury models. Mu opioid receptor agonists administered after injury lead to longer recovery times in models of spinal cord injury and neuropathic pain (^48–51^), while kappa opioid receptor agonists may be protective ^52^. In regard to TBI, the impact of opioid exposure on injury outcomes is less clear. Fentanyl administered immediately following controlled cortical impact exacerbated motor and spatial learning deficits within three weeks of injury ^53^. In a comparison of multiple anesthetic agents delivered immediately after controlled cortical impact, morphine was associated with the worst outcomes on motor performance in the first five days after injury, while fentanyl was associated with the worst performance in the Morris water maze two weeks following injury ^54^. Conversely, there is also evidence that opioids may be protective in experimental TBI. Morphine given immediately after injury was found to be beneficial in weight drop TBI-associated deficits in spatial learning ability 30-90 days post-injury ^55^. The mu opioid receptor selective agonist DAMGO delivered i.c.v. immediately prior to injury protected against beam walk and beam balance deficits in a rat fluid percussion injury model, and these outcomes were worsened by the mu antagonist beta-funaltrexamine ^56^. In the present study, we observed several interactive effects between rbTBI and oxycodone self-administration consistent with worse outcomes. In addition to injury exacerbating oxycodone seeking, injury x oxycodone self-administration interactions on FA were found in the NAc. Functional measures were more sensitive to interaction effects, with significant effects identified in the mPFC, NAc, and several other regions in local connectivity (REHO) and widespread functional connectivity. Furthermore, functional connectivity between the left and right IL mPFC positively correlated with both oxycodone seeking and intake in the combined analysis of sham and injured rats, and the left IL-NAc (ipsilateral to injury) connectivity positively correlated with drug intake in analysis of rbTBI rats alone and when combined with shams. These measurements we taken following measures of drug seeking – future studies will need to be done in order to determine if these metrics are predictive of future drug seeking.

### Neuroimaging studies of TBI

Diffusion tensor imaging (DTI) is a sensitive measure to identify microstructural damage associated with TBI (reviewed in ^57–59^). Experimental blast TBI has been shown to lead to widespread decreases in FA^13^, with closely-spaced repeated blast exposures exacerbating these effects ^60^. DTI measurements also tend to be exacerbate with time post-injury: we have reported an increase in the volume of microstructural damage between post-injury day 4 and 30 ^13^, and Badea et al. have reported greater changes in DTI at 90 days relative to 7 days post-injury ^61^. Another study found region-specific changes in DTI between 1 and 14 days after injury, reflecting regional differences in time to vasogenic or cytotoxic edema and demyelination ^62^. In the present study, we did not observe a significant effect of rbTBI on FA in the prefrontal cortex or nucleus accumbens, but we did observe a significant interaction between rbTBI and oxycodone self-administration in the nucleus accumbens. Despite a main or interactive effect in the PFC, we did observe correlations between FA and oxycodone seeking that were largely driven by animals that also experienced rbTBI.

Changes in rsfc have been observed in human TBI and in experimental TBI. Increased intra-hemispheric connectivity of the anterior cingulate cortex was found in veterans and athletes that had experienced a mild TBI ^63, 64^, an effect we observed in the PrL (immediately ventral of the anterior cingulate cortex in the rat). In collegiate athletes, elevated local connectivity (REHO) in the right middle and superior frontal gyri was associated with clinical symptoms following sports concussion ^65^. Widespread disruptions in functional connectivity have also been associated with long-term effects of TBI (reviewed in ^66–68^). In a study of high school and collegiate football players, increased widespread rsfc was observed only in athletes that remained symptomatic one week following injury ^69^. Similarly, elevated widespread rsfc was found in adolescent hockey players at three months following injury, although this effect was greater in recovered athletes relative to those with fewer symptoms ^70^. Combat veterans that sustained blast mild TBI and had persistent post-concussive symptoms had increased rsfc (assessed by magnetoencephalography) in several brain regions, including the ventromedial cortex and anterior cingulate cortices ^71^. Similarly, increases in functional connectivity in moderate and severe TBI were found for up to a year after injury, and the changes were greatest in areas that were already highly connected ^72^. Conversely, other studies using BOLD MRI have reported lower rsfc and/or local connectivity in injured patients ^73, 74^. In a study of TBI across different injury mechanisms, increased local connectivity was observed, and in a subset of patients, areas of highest connectivity were colocalized with high [^18^F]AV-1451 binding, a marker of tau hyperphosphorylation ^75^. The authors speculate that this may be associated with vascular neuropathology. Consistent with many of these studies, we found increased widespread and local connectivity. Similar to our observation of increased inter-hemispheric connectivity within the mPFC, Kulkarni et al. reported increased rsfc within PrL and IL after a single closed head momentum exchange injury ^76^.

### Neuroimaging studies of opioid use

Clinical studies have consistently identified structural and functional changes associated with opioid abuse. A voxel-based meta-analysis of studies employing DTI found that bilateral reductions in FA in frontal subgyral regions, including areas impacting the communication between cingulate and limbic association cortices with prefrontal association cortices ^77^. Heroin dependent individuals were also found to have lower FA in the white matter tract connecting the mPFC and postcentral gyrus ^78^ and a comparison of heroin users that relapsed to those that remained abstinent 6 months after initiation of methadone maintenance therapy found lower FA in several white matter tracts in the relapse group ^79^. In a neuroimaging study of over 150 subjects, increased widespread structural connectivity was reported in abstinent heroin users relative to non-heroin using controls ^80^. In this study, white matter FA was increased in heroin users, and FA within frontal networks was correlated with daily heroin intake ^80^.

Opioid use and abstinence have also been associated with altered functional neuroimaging measures in human populations. In a study of former heroin users undergoing methadone maintenance, those that relapsed had elevated REHO in medial orbital cortex and the right caudate nucleus, and elevated REHO in the caudate was correlated with relapse rates ^81^. In contrast, methadone maintained former heroin users had lower REHO in the medial orbital cortex compared to control subjects without a history of heroin or methadone use ^82^. In the present study, we found that rats receiving rbTBI and oxycodone self-administration had elevated REHO in several prefrontal regions, including medial and lateral orbital cortex. A trace of this effect was evident in the orbital cortex from blast alone, but we observed no indication of this in animals exposed to oxycodone only. Differences between our preclinical data and the aforementioned clinical data may arise from the interactive effects of the oxycodone and injury, the lack of opioid maintenance therapy in our rodent study, or other experimental differences. Zhou et al. found increased functional connectivity between the ventromedial PFC and nucleus accumbens in former heroin users with >= 3 years of abstinence ^83^, similar to findings of increased functional connectivity between ventral/rostral anterior cingulate cortex and the nucleus accumbens ^84^. Increased widespread functional connectivity has been also been reported in heroin use during abstinence ^85^, although there are also findings of decreased rsfc in opioid addiction, especially when examining connectivity within an executive network (reviewed in ^86, 87^). In conclusion, we find evidence that repeated mild TBI and oxycodone interact in such a way that manifests as elevated drug seeking, increased microstructural damage, and widespread aberrant functional connectivity. Repeated blast mild TBI also increased the effect sizes of many relationships between the neuroimaging findings and prior drug seeking and/or intake.

## Acknowledgements

This research was supported by National Institute on Drug Abuse Grant DA039276, United States Department of Veteran Affairs grant RX002931and the Research and Education Initiative Fund, a component of the Advancing a Healthier Wisconsin Endowment at the Medical College of Wisconsin. The content is solely the responsibility of the authors and does not necessarily represent the official views of the National Institutes of Health.

